# Modeling allele-specific gene expression by single-cell RNA sequencing

**DOI:** 10.1101/109629

**Authors:** Yuchao Jiang, Nancy R Zhang, Mingyao Li

## Abstract

Allele-specific expression is traditionally studied by bulk RNA sequencing, which measures average expression across cells. Single-cell RNA sequencing (scRNA-seq) allows the comparison of expression distribution between the two alleles of a diploid organism and thus the characterization of allele-specific bursting. We propose SCALE to analyze genome-wide allele-specific bursting, with adjustment of technical variability. SCALE detects genes exhibiting allelic differences in bursting parameters, and genes whose alleles burst non-independently. We apply SCALE to mouse blastocyst and human fibroblast cells and find that, globally, *cis* control in gene expression overwhelmingly manifests as differences in burst frequency.

## Background

In diploid organisms, two copies of each autosomal gene are available for transcription, and differences in gene expression level between the two alleles are widespread in tissues [1–7]. Allele-specific expression (ASE), in its extreme, is found in genomic imprinting, where the allele from one parent is uniformly silenced across cells, and in random X-chromosome inactivation, where one of the two X-chromosomes in females is randomly silenced. During the last decade, using single-nucleotide polymorphism (SNP)-sensitive microarrays and bulk RNA sequencing (RNA-seq), more subtle expression differences between the two alleles were found, mostly in the form of allelic imbalance of varying magnitudes in mean expression across cells [8–11]. In some cases such expression differences between alleles can lead to phenotypic consequences and result in disease [3, 12–14]. These studies, though revelatory, were at the bulk tissue level, where one could only observe average expression across a possibly heterogeneous mixture of cells.

Recent developments in single-cell RNA sequencing (scRNA-seq) have made possible the better characterization of the nature of allelic differences in gene expression across individual cells [6, 15, 16]. For example, recent scRNA-seq studies estimated that 12-24% of the expressed genes are monoallelically expressed during mouse preimplantation development [2] and that 76.4% of the heterozygous loci across all cells express only one allele [17]. These ongoing efforts have improved our understanding of gene regulation and enriched our vocabulary in describing gene expression at the allelic level with single-cell resolution.

Despite this rapid progress, much of the potential offered by scRNA-seq data remains untapped. ASE, in the setting of bulk RNA-seq data, is usually quantified by comparing the mean expression level of the two alleles. However, due to the inherent stochasticity of gene expression across cells, the characterization of ASE using scRNA-seq data should look beyond mean expression. A fundamental property of gene expression is transcriptional bursting, in which transcription from DNA to RNA occurs in bursts, depending on whether the gene’s promoter is activated (Figure 1A) [18, 19]. Transcriptional bursting is a widespread phenomenon that has been observed across many species including bacteria [20], yeast [21], Drosophila embryos [22], and mammalian cells [23, 24], and is one of the primary sources of expression variability in single cells. Figure 1B illustrates the expression across time of the two alleles of a gene. Under the assumption of ergodicity, each cell in a scRNA-seq sample pool is at a different time in this process, implying that for each allele, some cells might be in the transcriptional “ON” state, whereas other cells are in the “OFF” state. While in the “ON” state, the magnitude and length of the burst can also vary across cells, further complicating analysis. For each expressed heterozygous site, a scRNA-seq experiment gives us the bivariate distribution of the expression of its two alleles across cells, allowing us to compare the alleles not only in their mean, but also in their distribution. In this paper, we will use scRNA-seq data to characterize transcriptional bursting in an allele-specific manner and detect genes with allelic differences in the parameters of this process.

**Figure 1.**
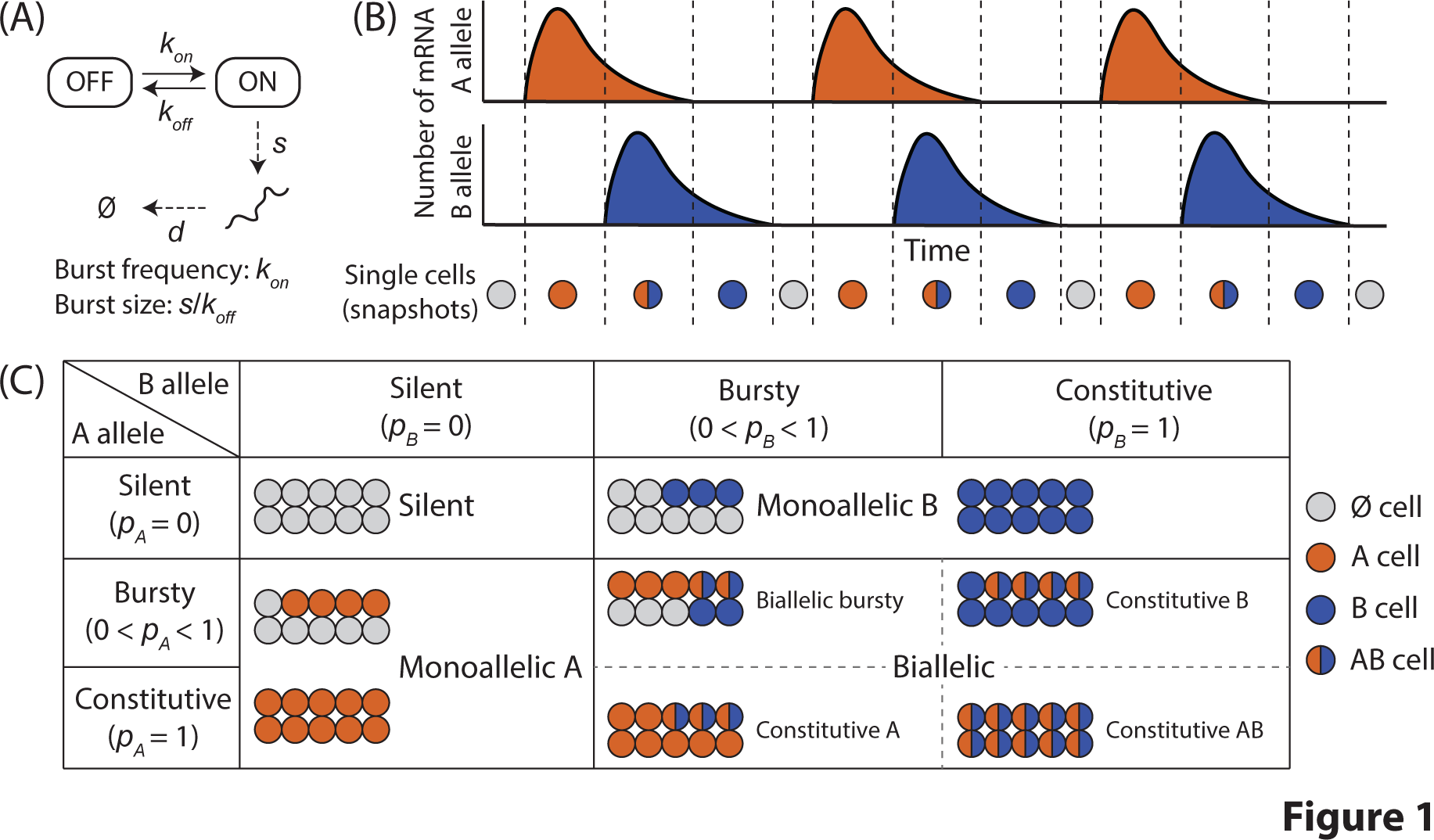
Allele-specific transcriptional bursting and gene categorization by single-cell ASE. (A) Transcription from DNA to RNA occurs in bursts, where genes switch between the “ON” and the “OFF” states. *k_on_*, *k_off_*, *s*, and *d* are activation, deactivation, transcription, and mRNA decay rate in the kinetic model respectively. (B) Transcriptional bursting of the two alleles of a gene give rise to cells expressing neither, one, or both alleles of a gene, sampled as vertical snapshots along the time axis. Partially adapted from Reinius and Sandberg [6]. (C) Empirical Bayes framework that categorizes each gene as silent, monoallelic and biallelic (biallelic bursty, one-allele constitutive, and both-alleles constitutive) based on ASE data with single-cell resolution.

Kim and Marioni [25] first studied bursting kinetics of stochastic gene expression from scRNA-seq data, using a Beta-Poisson model and estimated the kinetic parameters via a Gibbs sampler. In this early attempt, they assumed shared bursting kinetics between the two alleles and modeled total expression of a gene instead of allele-specific expression. Current scRNA-seq protocols often introduce substantial technical noise (Figure S1) [26–30], and these noise (e.g., gene dropouts, amplification and sequencing bias) are largely ignored in Kim and Marioni [25] and another recent scRNA-seq study Borel et al. [17], where, in particular, gene dropout may have led to overestimation of the pervasiveness of monoallelic expression (ME). Realizing this, Kim et al. [31] incorporated measurements of technical noise from external spike-in molecules into the identification of stochastic ASE (defined as excessive variability in allelic ratios among cells), and concluded that more than 80% of stochastic ASE in mouse embryonic stem cells are due to scRNA-seq technical noise. Kim et al.’s analysis was restricted to the identification of random monoallelic expression (RME) and did not consider more general patterns of ASE such as allele-specific transcriptional bursting.

ScRNA-seq also enables us to quantify the degree of dependence between the expressions of the two alleles. A previous RNA fluorescence *in situ* hybridization (FISH) experiment fluorescently labeled 20 genes in an allele-specific manner and showed that there was no significant deviation from independent bursting between the two alleles [32]. A recent scRNA-seq study of mouse cells through embryonic development [2] produced similar conclusions on the genome-wide level: They modeled transcript loss by splitting each cell’s lysate into two fractions of equal volume and controlling for false discoveries by diluting bulk RNA down to single-cell level. Their results suggest that on the genome-wide scale, assuming both alleles share the same bursting kinetics, the two alleles of most genes burst independently. Deviation from the theoretical curve in Deng et al. [2] for independent bursting with shared allele-specific kinetics, however, can be due to not only dependent bursting, but also differential bursting kinetics.

In this paper, we develop SCALE (**<u>S</u>**ingle-**<u>C</u>**ell **<u>AL</u>**lelic **<u>E</u>**xpression), a systematic statistical framework to study ASE in single cells by examining allele-specific transcriptional bursting kinetics. Our main goal is to detect and characterize differences between the two alleles in their expression distribution across cells. As a by-product, we will also quantify the degree of dependence between the expressions of the two alleles. SCALE is comprised of three steps. First, an empirical Bayes method determines, for each *gene*, whether it is silent, monoallelically expressed, or biallelically expressed, based on its allele-specific counts across cells (Figure 1C). Next, for genes determined to be biallelic bursty (i.e., both alleles have zero expression level in some but not all cells), a Poisson-Beta hierarchical model is used to estimate allele-specific transcriptional kinetics while accounting for technical noise and cell size differences. Finally, resampling-based testing procedures are developed to detect allelic differences in transcriptional burst size or burst frequency, and identify genes whose alleles exhibit non-independent transcription.

*In silico* simulations are conducted to investigate estimation accuracy and testing power. The stringency of model assumptions, and the robustness of the proposed procedures to the violation of these assumptions, will be discussed as they are introduced. Using SCALE, we reanalyze the scRNA-seq data for 122 mouse blastocyst cells [2] and 104 human fibroblast cells [17]. The mouse blastocyst study initially found abundant RME generated by independent and stochastic allelic transcription [2]; the human fibroblast study reported that 76.4% of the heterozygous loci displayed patterns of ME [17]. Through proper modeling of technical noise, our re-analysis of these two datasets brings forth new insights: While for 90% of the bursty genes, there are no significant deviations from the assumption of independent allelic bursting and shared bursting kinetics, the remaining bursty genes show differential burst frequency by a cis-effect and/or non-independent bursting with an enrichment in coordinated bursting. Collectively, we present a genome-wide approach to systematically analyze expression variation in an allele-specific manner with single-cell resolution. SCALE is an open-source R package available at https://github.com/yuchaojiang/SCALE.

## Results

### Methods overview

Figure 2 shows an overview of the analysis pipeline of SCALE. We start with allele-specific read counts of endogenous RNAs across all profiled single cells. An empirical Bayes method is adopted to classify expression of genes into monoallelic, biallelic, and silent states based on ASE data across cells. SCALE then estimates allele-specific transcriptional bursting parameters via a hierarchical Poisson-Beta model, while adjusting for technical variabilities and cell size differences. Statistical testing procedures are then performed to identify genes whose two alleles have different bursting parameters or burst non-independently. We describe each of these steps in turn.

**Figure 2.**
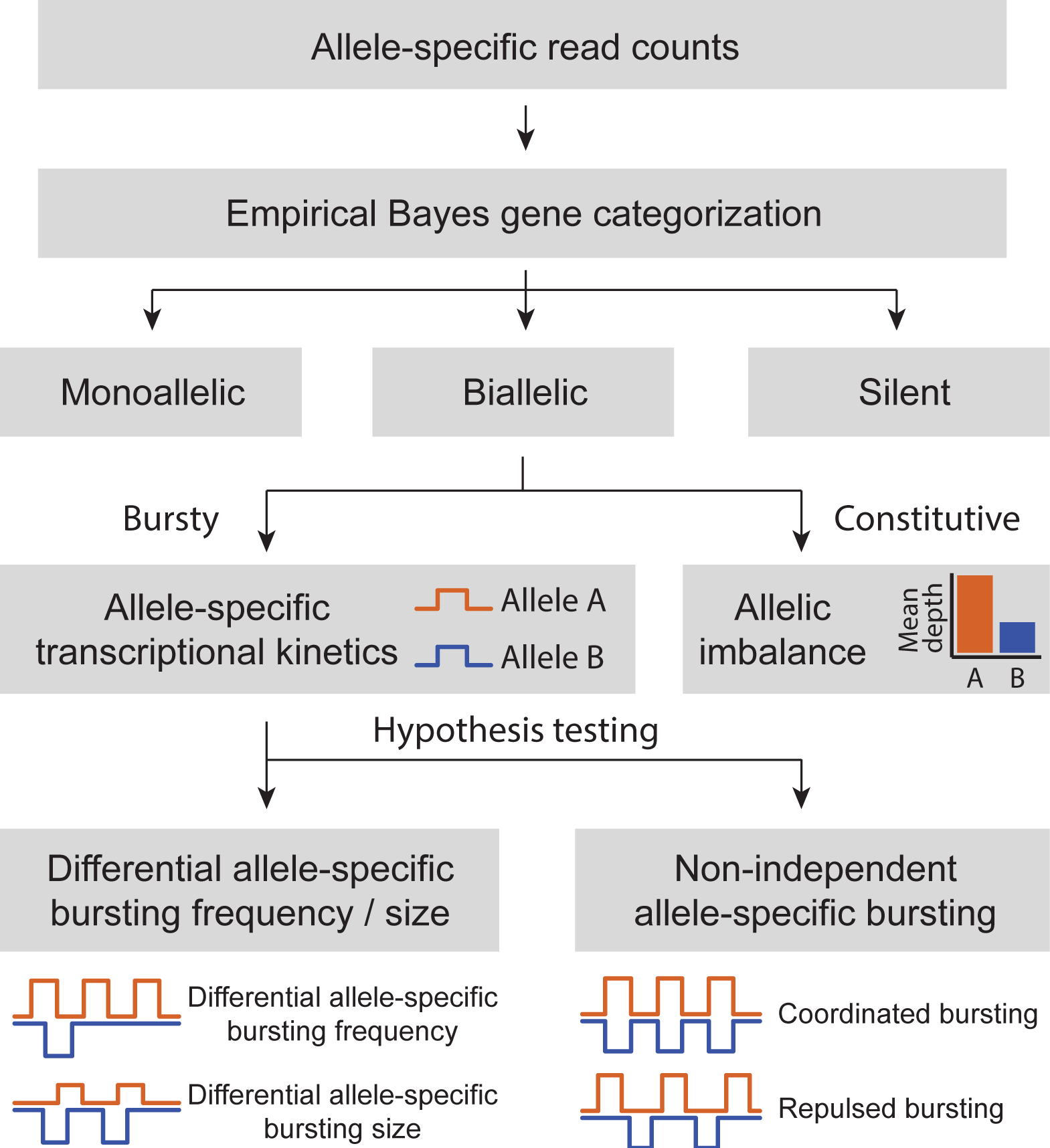
Overview of analysis pipeline of SCALE. SCALE takes as input allele-specific read counts at heterozygous loci and carries out three major steps: (i) an empirical Bayes method for gene classification, (ii) a Poisson-Beta hierarchical model to estimate allele-specific transcriptional kinetics with adjustment of technical variability and cell size, (iii) a hypothesis testing framework to test the two alleles of a gene have differential bursting kinetics and/or non-independent firing.

#### Gene classification by ASE data across cells

SCALE first determines for each gene whether its expression is silent, paternal/maternal monoallelic, or biallelic. Figure 1C outlines this categorization scheme. Briefly, for each gene, each cell is assigned to one of four categories corresponding to scenarios where both alleles are off (∅), only A allele is expressed (*A*), only B allele is expressed (*B*), and both alleles are expressed (*AB*). An expectation-maximization (EM) algorithm is implemented for parameter estimation. This classification accounts for both sequencing depth variation and sequencing errors. The assignment of the *gene* is then determined based on the posterior assignments of all cells. For example, if all cells are assigned to {∅}, the gene is silent; if all cells are assigned to either {∅} or {*A*}, the gene has ME of the A allele; if all cells are assigned to either {∅} or {*B*}, the gene has ME of the B allele; if both A and B allele are expressed in the cell pool, then the gene is biallelically expressed. Refer to Methods for detailed statistical method and the EM algorithm.

Through simulation studies (under section Assessment of estimation accuracy and testing power), we show that bursting parameters can only be stably estimated for *bursty* genes, that is, genes that are silent in a non-zero proportion of cells. Therefore, for biallelic bursty genes, allele-specific transcriptional kinetics are modeled through a Poisson-Beta distribution with adjustment of technical noise. For silent, monoallelically expressed, or constitutively expressed genes, there is no way nor need to estimate bursting kinetics for both alleles.

#### Allele-specific transcriptional bursting

When studying ASE in single cells, it is critical to consider transcriptional bursting due to its pervasiveness in various organisms [20–24]. We adopt a Poisson-Beta hierarchical model to quantify allele-specific transcriptional kinetics while accounting for dropout events and amplification and sequencing bias. Here, we start by reviewing the relevant literature with regard to transcriptional bursting at the single-cell level.

A two-state model for gene transcription is shown in Figure 1A, where genes switch between the “ON” and “OFF” states with activation and deactivation rates *k_on_* and *k_off_*. When the gene is at the “ON” state, DNA is transcribed into RNA at rate *s* while RNA decays at rate *d*. A Poisson-Beta stochastic model was firstly proposed by Kepler and Elston [33]:

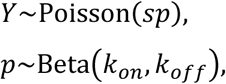

where *Y* is the number of mRNA molecules and *p* is the fraction of time that the gene spends in the active state, the latter having mean *k_on_*/(*k_on_* + *k_off_*). Under this model, 1/*k_on_* and 1*k*/_*off*_ are the average waiting times in the inactive and active states, respectively. *Burst size,* defined as the average number of synthesized mRNA per burst episode, is given by *s*/*k*_*off*_, and *burst frequency* is given by *k_on_*. Kepler and Elston [33] gave detailed analytic solutions via differential equations. Raj et al. [23] offered empirical support for this model via single-molecule FISH experiment on reporter genes. Since the kinetic parameters are measured in units of time and only the stationary distribution is assumed to be observed (e.g., when cells are killed for sequencing and fixed for FISH experiment), the rate of decay *d* is set to one [15]. This is equivalent to having three kinetic parameters {*s, k_on_, k_off_*}, each normalized by the decay rate *d*. Kim and Marioni [25] applied this Poisson-Beta model to total gene-level transcript counts from scRNA-seq data of mouse embryonic stem cells. While they found that the inferred kinetic parameters are correlated with RNA polymerase II occupancy and histone modification [25], they didn’t address the issue of technical noise, especially the dropout events, introduced by scRNA-seq. Failure of accounting for gene dropouts may lead to biased estimation of bursting kinetics.

Furthermore, since the transitions between active and inactive states occur separately for the two alleles, when allele-specific expression data are available, it seems more appropriate to model transcriptional bursting in an allele-specific manner. The fact that transcriptional bursting occurs independently for the two alleles has been supported by empirical evidence: Case studies based on imaging methods have suggested that the two alleles of genes are transcribed in an independent fashion [34, 35]; using scRNA-seq data, Deng et al. [2] showed that the two alleles of most genes tend to fire independently with the assumption that both alleles share the same set of kinetic parameters. These findings, although limited in scale or relying on strong assumptions, emphasize the need to study transcriptional bursting in an allele-specific manner.

#### Technical noise in scRNA-seq and other complicating factors

Figure S1 outlines the major steps of the scRNA-seq protocols and the sources of bias that are introduced during library preparation and sequencing. After the cells are captured and lysed, exogenous spike-ins are added as internal controls, which have fixed and known concentration and can thus be used to convert the number of sequenced transcripts into actual abundances. During the reverse transcription, pre-amplification, and library preparation steps, lowly expressed transcripts might be lost, in which case they will not be detected during sequencing. This leads to the so-called “dropout” events. Since spike-ins undergo the same experimental procedure as endogenous RNAs in a cell, amplification and sequencing bias can be captured and estimated through the spike-in molecules. Here we adopt the statistical model in TASC (Toolkit for Analysis of Single Cell data, unpublished), which explicitly models the technical noise through spike-ins. TASC’s model is based on the key observation that the probability of a gene being a “dropout” depends on its true expression in the cell, with lowly expressed gene more likely to drop out. Specifically, let *Q_cg_* and *Y_cg_* be, respectively, the observed and true expression level of gene *g* in cell *c*. The hierarchical mixture model used to model dropout, amplification and sequencing bias is:

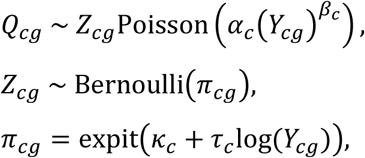

where *Z*_*cg*_ is a Bernoulli random variable indicating that gene *g* is detected in cell *c*, that is, a dropout event has not occurred. The success probability *π_cg_=P(Z_cg_=1)* depends on log(*Y_cg_*), the logarithm of the true underlying expression. Cell-specific parameters *α_c_* models the capture and sequencing efficiency; *β_c_* models the amplification bias; *κ_c_* and *τ_c_* characterize whether a transcript is successfully captured in the library. This model will later be used to adjust for technical noise in allele-specific expression.

As input to SCALE, we recommend scRNA-seq data from cells of the same type. Unwanted heterogeneity, however, still persists as the cells may differ in size or may be in different phases of the cell cycle. Through a series of single-cell FISH experiments, Padovan-Merhar et al. [36] showed how gene transcription depends on these exogenous factors: burst size is independent of cell cycle but is kept proportional to cell size by a *trans* mechanism; burst frequency is independent of cell size but is reduced approximately by half, through a *cis* mechanism, between G1 and G2 phase to compensate for the doubling of DNA content. Figure S2 gives an illustration on how burst size and burst frequency change with cell size and cell cycle phase. Note that, while the burst frequency from *each* DNA copy is halved when the amount of DNA is doubled, the total burst frequency remains roughly constant through the cell cycle. Thus, SCALE adjusts for variation in cell size through modulation of burst size, and does not adjust for variation in cell cycle phase. Details will be given below.

There are multiple ways to measure cell size. Padovan-Merhar et al. [36] proposed using the expression level of *GAPDH* as a cell size marker. When spike-ins are available, we use the ratio of the total number of endogenous RNA reads over the total number of spike-in reads as a measure (Figure S2) of the total RNA volume, which was shown to be a good proxy for cell size [28]. SCALE allows the user to input the cell sizes *ϕ_c_*, if these are available through other means.

#### Modeling transcriptional bursting with adjustment of technical and cell-size variation

We are now ready to formulate the allele-specific bursting model for scRNA-seq data. For genes that are categorized as biallelic bursty (with proportion of cells expressing each allele between 5% and 95% from the Bayes framework), SCALE proceeds to estimate the allele-specific bursting parameters using a hierarchical model:

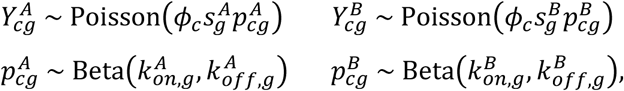

where 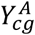 and 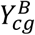 are the true allele-specific expressions for gene *g* in cell *c*. The two alleles of each gene are modeled by separate Poisson-Beta distributions with kinetic parameters that are gene- and allele-specific. These two Poisson-Beta distributions share the same cell size factor *ϕ*_*c*_, which affects burst size. The true allele-specific expressions 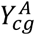 and 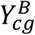 are not directly observable. The observed allele-specific read counts 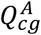 and 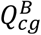 are confounded with technical noise, and follow the Poisson mixture model outlined in the previous section:

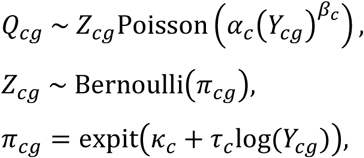

How to generate input to SCALE for both endogenous RNAs and exogenous spike-ins is included in Methods and Supplementary Methods. For parameter estimation, we developed a new “histogram-repiling” method to obtain the distribution of *Y_cg_* from the observed distribution of *Q_cg_*. The bursting parameters are then derived from the distribution of *Y_cg_* by moment estimators. Standard errors and confidence intervals of the parameters are obtained using nonparametric bootstrap. The details are shown in Methods.

#### Hypothesis testing

For biallelic bursty genes, we use nonparametric Bootstrap to test the null hypothesis that the burst frequency and burst size of the two alleles are the same 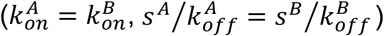 against the alternative hypothesis that either or both parameters differ between alleles. For each gene, we also perform chi-square test to determine if the transcription of the two alleles are independent by comparing the observed proportions of cells from the gene categorization framework against the expected proportions under independence. For genes where the proportion of cells expressing both alleles is significantly higher than expected, we define their bursting as coordinated; for genes where the proportion of cells expressing only one allele is significantly higher than expected, we define their bursting as repulsed (Figure 2). We adopt false discovery rate (FDR) to adjust for multiple comparisons. Details of the testing procedures are outlined in Methods.

### Analysis of scRNA-seq dataset of mouse cells during preimplantation development

We re-analyze the scRNA-seq dataset of mouse blastocyst cells dissociated from *in vivo* F1 embryos (CAST/female x C57/male) from Deng et al. [2]. Transcriptomic profiles of each individual cell was generated using the Smart-seq [37] protocol. For 22,958 genes, reads per kilo base per million reads (RPKM) and total number of read counts across all cells are available. Parental allele-specific read counts are also available at heterozygous loci (Figure S3). Principal component analysis (PCA) was performed on cells from oocyte to blastocyst stages of mouse preimplantation development and showed that the first three principal components well separate the early-stage cells from the blastocyst cells (Figure S4). The cluster of early-, mid-, and late-blastocyst cells are combined to gain sufficient sample size. In discussion, we give further insights on the potential effects of cell subtype confounding. Quality control (QC) procedure was adopted to remove outliers in library size, mean and standard deviation of allelic read counts/proportions. We apply SCALE to this dataset of 122 mouse blastocyst cells, with a focus on addressing the issue of technical variability and modeling of transcriptional bursting.

Eight exogenous RNAs with known serial dilutions are added in late blastocyst cells (Table S1A) and are used to estimate the technical-noise associated parameters (Figure S5A). We apply the Bayes gene classification framework to these cells to get the genome-wide distribution of gene categories. Specifically, out of the 22,958 genes profiled across all cells, ~43% are biallelically expressed (~33% of the total are biallelic bursty and ~10% of the total are biallelic non-bursty), ~7% are monoallelically expressed, and ~50% are silent. Our empirical Bayes categorization results show that, on the genome-wide scale, the two alleles of most biallelic bursty genes share the same bursting kinetics and burst independently (Figure S6A), as has been reported by Deng et al. [2].

For the 7,486 genes that are categorized as biallelic bursty, we apply SCALE to identify genes whose alleles have different bursting kinetic parameters by the Bootstrap-based hypothesis tests as previously described. After FDR control, we identify 425 genes whose two alleles have significant differential burst frequency (Figure 3A) and 2 genes whose two alleles have significant differential burst size (Figure 3B). Figure 4 shows the allelic read counts of a gene that has differential burst frequency (*Btf3l4*) and a gene that has differential burst size (*Fdps*). The two genes with significant differential allelic burst size, namely, gene *Fdps* and *Atp6ap2*, are also significant in having differential burst frequency between the two alleles. *P*-values from differential burst frequency testing have a spike below the significance level after FDR control (Figure 3A), while those from differential burst size testing are roughly uniformly distributed (Figure 3B).

**Figure 3.**
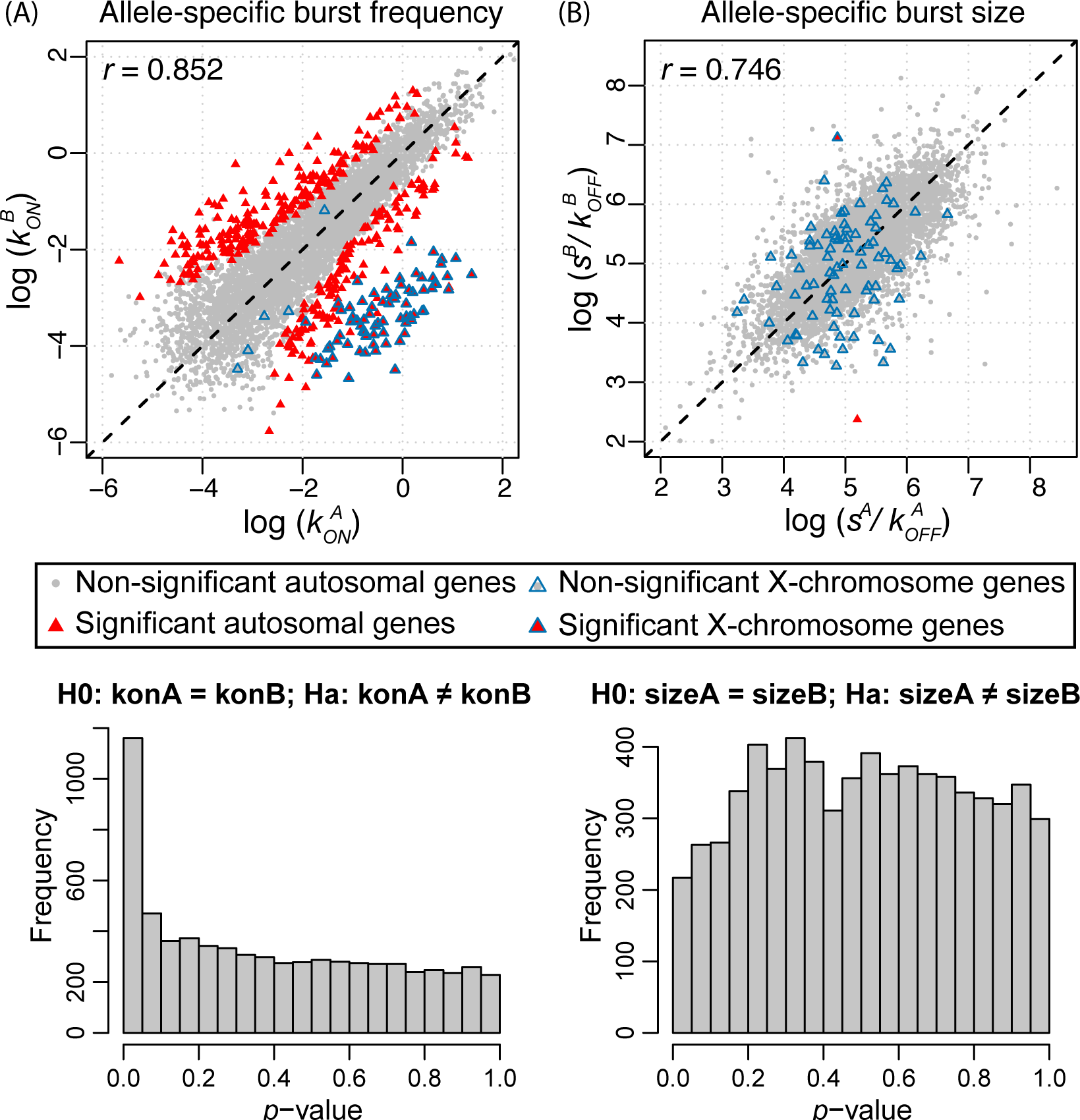
Allele-specific transcriptional kinetics of 7486 genes from 122 mouse blastocyst cells. (A) Burst frequency of the two alleles has a correlation of 0.852. 425 genes show significant allelic difference in burst frequency after FDR control. (B) Burst size of the two alleles has a correlation of 0.746. Two genes show significant allelic difference in burst size. X-chromosome genes as positive controls show significant higher burst frequencies of the maternal alleles than those of the paternal alleles. The *p*-values for allelic burst size difference (bottom right panels) are uniformly distributed as expected under the null, whereas those for allelic burst frequency difference (bottom left panels) have a spike below significance level after FDR control.

At the whole genome level, these results show that allelic differences in the expression of bursty genes during embryo development is achieved through differential modulation of burst frequency rather than burst size. This seems to agree with intuition, since allelic differences must be caused by factors that act in *cis* to regulate gene expression, and *cis* factors are likely to change burst frequency by affecting promoter accessibility [36, 38–39,40]. On the contrary, while it is plausible for *cis* factors to affect allelic burst size through, for example, the efficiency of RNA Polymerase II recruitment or the speed of elongation, the few known cases of burst size modulation are controlled in *trans* [36]. Furthermore, previous studies have shown that the kinetic parameter that varies the most – along the cell cycle [36], between different genes [41], between different growth conditions [42], or under regulation by a transcription factor [43] – is the probabilistic rate of switching to the active state *k_on_*, while the rates of gene inactivation *K_off_* and of transcription *s* vary much less.

Our analysis includes 107 male cells (X^A^Y) and 15 female cells (X^A^X^B^) and this allows us to use those bursty X-chromosome genes as positive controls. As a result of this gender mixture, there are more cells expressing the maternal X^A^ allele compared to the paternal X^B^ allele. As shown in Figure 3, SCALE successfully detects these bursty X-chromosome genes with significant difference in allelic burst frequency but not in allelic burst size. If we only keep the 107 male cells, these X-chromosome genes are correctly categorized as monoallelically expressed – the bursting kinetics for the paternal X^B^ allele are not estimable – and in this case there is no longer a cluster of significant X-chromosome genes separated from the autosomal genes (Figure S8).

For biallelic bursty genes, we also used a simple Binomial test to determine if the mean allelic coverage across cells is biased towards either allele. This is comparable to existing tests of allelic imbalance in bulk tissue, although the total coverage across cells in this dataset is much higher than standard bulk tissue RNA-seq data. After multiple hypothesis testing correction, we identify 417 genes with significant allelic imbalance, out of which 238 overlap with the significant genes from the testing of differential bursting kinetics (Figure 5A). Inspection of the estimated bursting kinetic parameters in Figure 5A shows that, when the burst size and burst frequency of the two alleles change in the same direction (e.g., gene *Gprc5a* in Figure 5B), testing of allelic imbalance can detect more significant genes with higher power. This is not unexpected – a small insignificant increase in burst size adds on top of an insignificant increase in burst frequency resulting in a significant increase in overall expression levels between the two alleles. However, for genes in red in the top left and bottom right quadrants of Figure 5A, the test for differential bursting kinetics detects more genes than the allelic imbalance test. This is due to the fact that when burst size and burst frequency change in opposite directions (e.g., gene *Dhrs7* in Figure 5B), their effects cancel out when looking at the mean expression. Furthermore, even when the burst size does not change, if the change in burst frequency is small, by using a more specific model SCALE has higher power to detect it as compared to an analysis based on mean allelic imbalance. Overall, the allelic imbalance test and differential bursting test report overlapping but substantially different set of genes, with each test having its benefits. Compared to the allelic imbalance test, SCALE gives more detailed characterization of the nature of the difference by attributing the change in mean expression to a change in the burst frequency and/or burst size.

It is also noticeable that in Figure 5A the vertical axis, ∆*freq*, has a 50% wider range than the horizontal axis, ∆*size*. Therefore, while it is visually not obvious from this scatter plot, there are much more genes with large absolute ∆*freq* than with large absolute ∆*size*. Although the standard errors of these estimated differences are not reflected in the plot, given our testing results, those genes with large estimated differences in ∆*size* also have large standard errors in their estimates, which is further confirmed via simulations.

**Figure 4.**
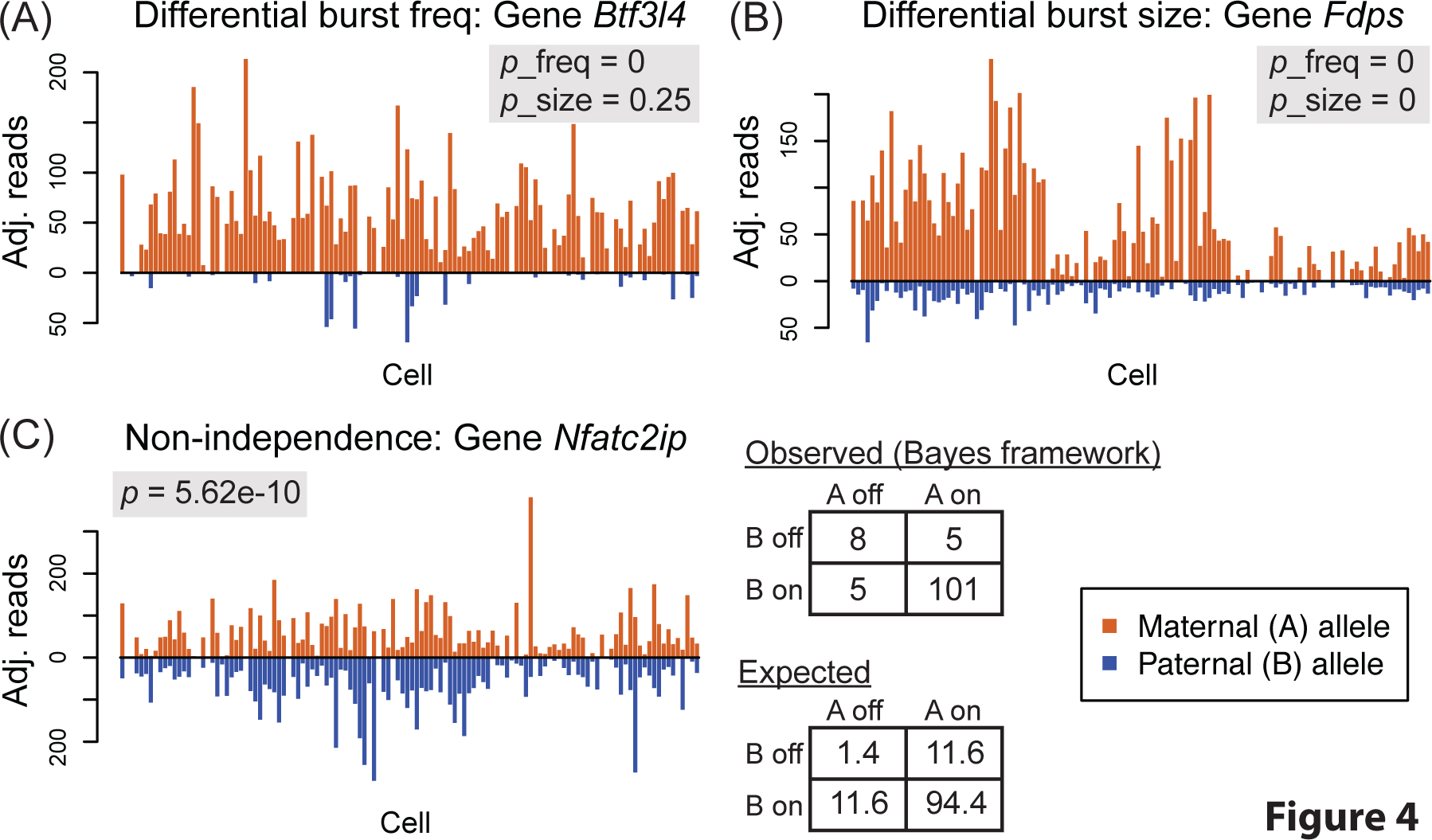
Examples of significant genes from hypothesis testing. (A) The two alleles of the gene have significantly differential burst frequency from the bootstrap-based testing. (B) The two alleles of the gene have significantly differential burst size and burst frequency. (C) The two alleles of the gene fire non-independently from the chi-square test of independence.

**Figure 5.**
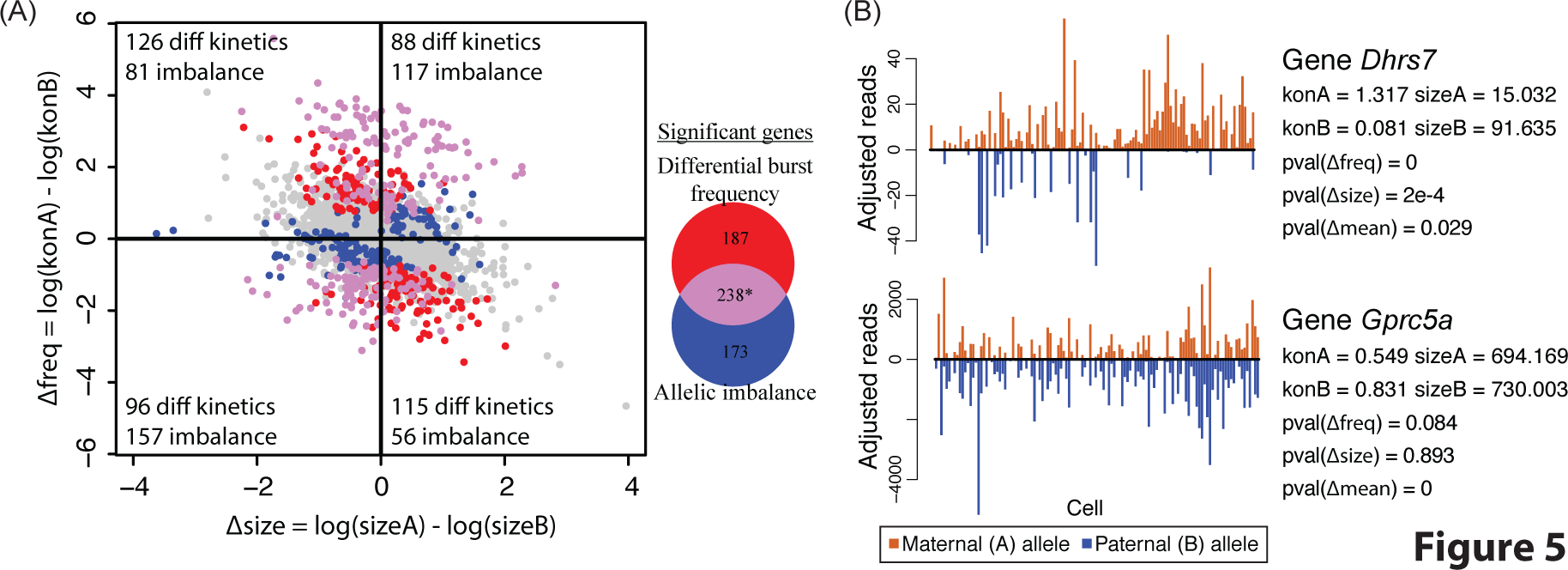
Testing of bursting kinetics by scRNA-seq and testing mean difference by bulk-tissue sequencing. (A) Venn diagram of genes that are significant from testing of shared burst frequency and allelic imbalance. *Also includes the two genes that are significant from testing of shared burst size. Change in burst frequency and burst size inthe same direction leads to higher detection power of allelic imbalance; change in different direction leads to allelic imbalance testing being underpowered. (B) Gene *Dhrs7* whose two alleles have bursting kinetics in different direction and gene *Gprc5a* whose two alleles have bursting kinetics in the same direction. *Dhrs7* is significant from testing of differential allelic bursting kinetics; *Gprc5a* is significant from the testing of mean difference between the two alleles.

Further chi-squared test of the null hypothesis of independence (Figure 4C) shows that there are 424 genes whose two alleles fire in a significantly non-independent fashion. We find that all significant genes have higher proportions of cells expressing both alleles than expected, indicating coordinated expression between the two alleles. In this dataset, there are no significant genes with repulsed bursting between the two alleles. Repulsed bursting, in the extreme case where at most one allele is expressed in any cell, is also referred to as stochastic ME [31]. Our testing results indicate that, in mouse embryo development, all cases of stochastic ME (i.e., repulsion between the two alleles) can be explained by independent and infrequent stochastic bursting. The burst synchronization in the 424 significant genes is not unexpected and is possibly due to a shared *trans* factor between the two alleles (e.g., co-activation of both alleles by a shared enhancer). This result is concordant with the findings from a mouse embryonic stem cell scRNA-seq study by Kim et al. [31], which reported that the two alleles of a gene show correlated allelic expression across cells more often than expected by chance, potentially suggesting regulation by extrinsic factors [31]. We further discuss the sharing of such extrinsic factors under the context of cell population admixtures in Discussion.

In summary, our results by SCALE suggest that: (i) The two alleles from 10% of the bursty genes show either significant deviations from independent firing or significant differences in bursting kinetic parameters, (ii) For genes whose alleles differ in their bursting kinetic parameters, the difference is found mostly in the burst frequency instead of the burst size, (iii) For genes whose alleles violate independence, their expression tends to be coordinated. Refer to Table S1B for genome-wide output from SCALE.

### Analysis of scRNA-seq dataset of human fibroblast cells

To further examine our findings in a dataset without potential confounding of cell type admixtures, we apply SCALE to a scRNA-seq dataset of 104 cells from female human newborn primary fibroblast culture from Borel et al. [17]. The cells were captured by Fluidigm C1 with 22 PCR cycles and were sequenced with on average 36 million reads (100 bp, paired end) per cell. Bulk-tissue whole genome sequencing was performed on two different lanes with 26-fold coverage on average and was used to identify heterozygous loci in coding regions. After QC procedures, 9016 heterozygous loci from 9016 genes were identified (if multiple loci coexist in the same gene, we pick the one with the highest mean depth of coverage). At each locus, we use SAMtools [44] mpileup to obtain allelic read counts in each single cell from scRNA-seq, which are further used as input for SCALE. 92 ERCC synthesized RNAs were added in the lysis buffer of 12 fibroblast cells with a final dilution of 1:40000. The true concentrations and the observed number of reads for all spike-ins are used as baselines to estimate technical variability (Table S1C, Figure S5B). Refer to Supplementary Methods for details on the bioinformatic pipeline.

We apply the gene categorization framework by SCALE and find that out of the 9016 genes, the proportions of monoallelically expressed, biallelically expressed, and silent genes are 11.5%, 45.7%, and 42.8%, respectively. For the 2277 genes that are categorized as biallelic bursty, we estimate their allele-specific bursting kinetic parameters and find that the correlations between the estimated burst frequency and burst size between the two alleles are 0.859 and 0.692 (Figure 6). We then carry out hypothesis testing on differential allelic bursting kinetics. After FDR correction, we identified 26 genes with significant differential burst frequency between the two alleles (Figure 6A) and one gene *Nfx1* with significantly differential burst size between the two alleles, which is also significant in burst frequency testing (Figure 6B). We further carry out testing of non-independent bursting between the two alleles and identify 35 significant genes after FDR correction (Figure S6B). Out of the 35 significant genes, 27 showed patterns of coordinated bursting while the rest 8 showed repulsed patterns. Refer to Table S1D for detailed output from SCALE across all tested genes.

**Figure 6.**
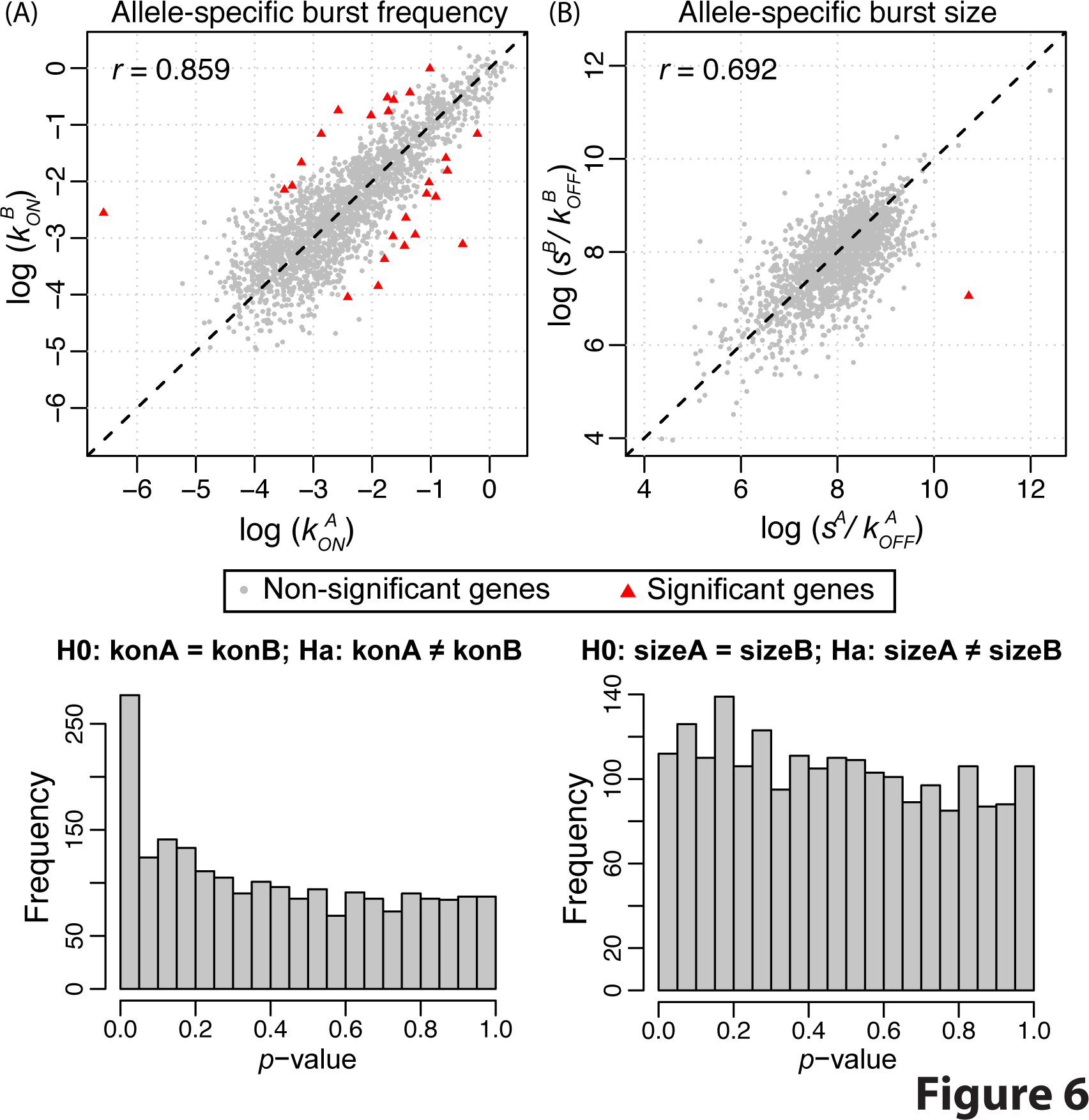
Allele-specific transcriptional kinetics of 2277 genes from 104 human fibroblast cells. (A) Burst frequency of the two alleles has a correlation of 0.859. 26 genes show significant allelic difference in burst frequency after FDR. (B) Burst size of the two alleles has a correlation of 0.692. One gene has significant allelic difference in burst size. The results are concordant with the findings from the mouse embryonic development study.

We also carry out pairwise correlation analysis between the estimated allelic bursting kinetics, the proportion of unit time that the gene stays in the active state *k_on_/(k_on_ + k_off_)* for each allele, as well as the overall allele-specific expression levels (taken as the sum across all cells at the heterozygous locus). Notably, we find that the overall allele-specific expression correlates strongly with the burst frequency and the proportion of time that the gene stays active, but not with the burst size (Figure S9), in concordance with Kim and Marioni [25]. This further supports our previous conclusion that the allele-specific expression at single-cell level manifests as differences in burst frequency in a *cis*-manner.

### Assessment of estimation accuracy and testing power

First, we investigate the accuracy of the moment estimators for the bursting parameters under four different scenarios in the Poisson-Beta transcription model: (i) small *k_on_* and small *k_off_*, which we call bursty and leads to relatively few transitions between the “ON” and “OFF” state with a bimodal mRNA distribution across cells (Figure S10A); (ii) large *k_on_* and small *k_off_*, which leads to long durations in the “ON” state and resembling constitutive expression with the mRNA having a Poisson-like distribution (Figure S10B); (iii) small *k_on_* and large *k_off_*, which leads to most cells being silent (Figure S10C); (iv) and large *k_on_* and large *k_off_*, which leads to constitutive expression (Figure S10D).

We generate simulated data for 100 cells from the four cases above and start with no technical noise or cell size confounding. Within each case, we vary *k_on_*, *k_off_* and *s* and use relative absolute error 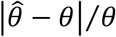 as a measurement of accuracy (Figure S11). Our results show that genes with large *k_on_* and small *k_off_* (shown as the black curves in Figure S11) have the largest estimation errors of the bursting parameters. Statistically it is hard to distinguish these constitutively expressed genes from genes with large *k_on_* and *k_off_* large and thus the kinetic parameters in this case cannot be accurately estimated, which has been previously reported [25, 45]. Furthermore, the estimation errors are large for genes with small *k_on_*, large *k_off_*, and small *s* (shown as red curves in Figure S11) due to lack of cells with nonzero expression. The standard errors and confidence intervals of the estimated kinetics from bootstrap resampling further confirm the underperformance for the above two classes (Table S2). This emphasizes the need to adopt the Bayes categorization framework as a first step so that kinetic parameters are stably estimated only for genes whose both alleles are bursty. For genes whose alleles are perpetually silent or constitutively expressed across cells, there is no good method, nor any need, to estimate their bursting parameters.

Importantly, we see that the estimation bias in transcription rate *s* and deactivation rate *k_off_* cancel – over/under estimation of *s* is compensated by over/under estimation of *k_off_* – and as a consequence the burst size *s*/*k_off_* can be more stably estimated than either parameter alone, especially when *k_on_* ≪ *k_off_* (shown as red curves in Figure S11). This is further confirmed by empirical results that allelic burst size has much higher correlation (0.746 from the mouse blastocyst dataset and 0.692 from the human fibroblast dataset) than allelic transcription and deactivation rate (0.464 and 0.265 for mouse blastocyst, and 0.458 and 0.33 for human fibroblast) (Figure S12). For this reason, all of our results on real data are based on *s*/*k_off_* and we do not consider *s* and *k_off_* separately.

We further carry out power analysis on the testing of differential burst frequency and burst size between the two alleles. The null hypothesis is both alleles sharing the same bursting kinetics 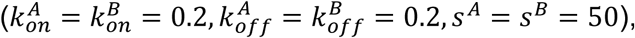 while the alternative hypotheses with differential burst frequency or burst size are shown in the legends in Figure S13. The detailed setup of the simulation procedures are as follows. (i) Simulate the true allele-specific read counts *Y^A^* and *Y^B^* across 100 cells from the Poisson-Beta model under the alternative hypothesis. Technical noise is then added based on the noise model described earlier with technical noise parameters {*α, β, κ, τ*} estimated from the mouse blastocyst cell dataset. (ii) Apply SCALE to the observed expression level *Q^A^* and *Q^B^*, which returns *p*-value for testing differential burst size or burst frequency. If the *p*-value is less than the significance level, we reject the null hypothesis. (iii) Repeat (i) and (ii) *N* times with the power estimated as
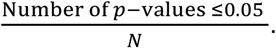 Our results indicate that the testing of burst frequency and burst size have similar power overall with relatively reduced power if the difference in allelic burst size is due to difference in the deactivation rate *k_off_*.

We then simulate allele-specific counts from the full model including technical noise as well as variations in cell size with the ground truth 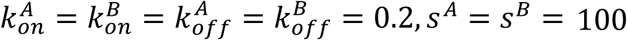 (bursty with small activation and deactivation rate). For parameters quantifying the degree of technical noise, we use the estimates from the mouse blastocyst cells (Figure S5A) as well as the human fibroblast cells (Figure S5B). Cell sizes are simulated from a normal distribution with mean 0 and standard deviation 0.1 and 0.01. We run SCALE under four different settings: (i) in its default setting, (ii) without accounting for cell size, (iii) without adjusting for technical variability, (iv) not in an allele-specific fashion but using total coverage as input. Each is repeated 5000 times with a sample size of 100 and 400 cells, respectively. Relative estimation errors of burst size and burst frequency are summarized across all simulation runs. Our results show that SCALE in its default setting has the smallest estimation errors for both burst size and burst frequency (Figure S14-S15). Not surprisingly, cell size has larger effect on burst size estimation than burst frequency estimation, while technical variability leads to biased estimation of both burst frequency and burst size. The estimates taking total expression instead of ASE as input are completely off. Furthermore, the estimation accuracy improved as the number of cells increased. These results indicate the necessity to profile transcriptional kinetics in an allele-specific fashion with adjustment of technical variability and cell size.

## Discussion

We propose SCALE, a statistical framework to study ASE using scRNA-seq data. The input data to SCALE are allele-specific read counts at heterozygous loci across all cells. In the two datasets that we analyzed, we use the F1 mouse crossing and the bulk-tissue sequencing to profile the true heterozygous loci. When these are not available, scRNA-seq itself can be used to retrieve allele-specific expression and more specifically haplotype, as illustrated in Edsgard et al. [46]. SCALE estimates parameters that characterize allele-specific transcriptional bursting, after accounting for technical biases in scRNA-seq and size differences between cells. This allows us to detect genes that exhibit allelic differences in burst frequency and burst size, and genes whose alleles show coordinated or repulsed bursting patterns. Differences in mean expression between the two alleles have long been observed in bulk RNA-seq. By scRNA-seq, we now move beyond the mean and characterize the difference in expression distributions between the two alleles, specifically in terms of their transcriptional bursting parameters.

Transcriptional bursting is a fundamental property of gene expression, yet its global patterns in the genome has not been well characterized, and most studies consider bursting at the gene level by ignoring the allelic origin of transcription. In this paper, we reanalyzed the Deng et al. [2] and Borel et al. [17] data. We confirmed the findings from Levesque and Raj [32] and Deng et al. [2] that for most genes across the genome there is no sufficient evidence against the assumption of independent bursting with shared bursting kinetics between the two alleles. For genes where significant deviations are observed, SCALE allows us to attribute the deviation to differential bursting kinetics and/or non-independent bursting between the two alleles.

More specifically, for genes that are transcribed in a “bursty” fashion, we compared the burst frequency and burst size, between their two alleles. For both scRNA-seq datasets, we identify significant number of genes whose allele-specific burstings differ in the burst frequency but not in the burst size. Our findings provide evidence that burst frequency, which represents the rate of gene activation, is modified in *cis*, and that burst size, which represents the ratio of transcription rate to gene inactivation rate, is less likely to be modulated in *cis*. Although our testing framework may have slightly reduced power in detecting differential deactivation rate (Figure S13), the regulation in burst size can either result from a global *trans* factor or extrinsic factors that acts upon both alleles. Similar findings have been previously reported, from different perspectives and on different scales, using various technologies, platforms, and model organisms [31, 36, 41–43].

It is worth noting that the estimated bursting parameters by SCALE are normalized by the decay rate, where the inverse 1/*d* denotes the average life time of an mRNA molecule. Here we implicitly make the assumptions that for each allele, the gene-specific decay rates (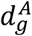 and 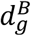) are constant, and thus the estimated allelic burst frequencies are the ratio of true burst frequency over decay rate (that is 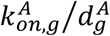 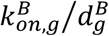 The decay rates, however, cancel out in the numerator and denominator in the allelic burst sizes, 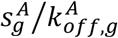 and 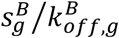 Therefore, the differences that we observe in the allelic burst frequencies can also potentially be due to differential decay rates between the two alleles, which has been previously reported to be regulated by microRNAs [47].

It is also important to note that 44% of the genes found to be significant for differential burst frequency are not significant in the allelic imbalance test based on mean expression across cells. This suggests that expression quantitative trait loci (eQTL) affecting gene expression through modulation of bursting kinetics is likely to escape detection in existing eQTL studies by bulk sequencing, especially when burst size and burst frequency change in different directions. This is further underscored by the study of Wills et al. [48], which measured the expression of 92 genes affected by Wnt signaling in 1,440 single cells from 15 individuals, and then correlated SNPs with various gene-expression phenotypes. They found bursting kinetics as characterized by burst size and burst frequency to be heritable, thus suggesting the existence of bursting-QTLs. Taken together, these results should further motivate more large scale genome-wide studies to systematically characterize the impact of eQTLs on various aspects of transcriptional bursting.

Kim et al. [31] described a statistical framework to quantify the extent of stochastic ASE in scRNA-seq data by using of spike-ins, where stochastic ASE is defined as excessive variability in the ratio of the expression level of the paternal (or maternal) allele between cells after controlling for mean allelic expression levels. While they attributed 18% of the stochastic ASE to biological variability, they did not examine what biological factors lead to these stochastic ASE. In this paper, we attribute the observed stochastic ASE to difference in allelic bursting kinetics. By studying bursting kinetics in an allele-specific manner, we can compare the transcriptional differences between the two alleles at a finer scale.

Kim and Marioni [25] described a procedure to estimate bursting kinetic parameters using scRNA-seq data. Our method differs from Kim and Marioni [25] in several ways. First, our model is an allele-specific model that infers kinetic parameters for each allele separately, thus allowing comparisons between alleles. Second, we infer kinetic parameters based on the distribution of “true expression” rather than the distribution of observed expression. We are able to do this through the use of a simple and novel deconvolution approach, which allows us to eliminate the impact of technical noise when making inference on the kinetic parameters. Appropriate modeling of technical noise, in particular, gene dropouts, is critical in this context, as failing to do so could lead to the overestimation of *k_off_*. Third, we employ a gene categorization procedure prior to fitting the bursting model. This is important because the bursting parameters can only be reliably estimated for genes that have sufficient expression and that are bursty.

As a by-product, SCALE also allows us to rigorously test, for scRNA-seq data, whether the paternal and maternal alleles of a gene are independently expressed. In both scRNA-seq datasets we analyzed, we identified more genes whose allele-specific burstings are in a coordinated fashion than those in a repulsed fashion. The tendency towards coordination is not surprising, since the two alleles of a gene share the same nuclear environment and thus the same ensemble of transcription factors. We are aware that this degree of coordination can also arise from the mixture of non-homogeneous cell populations, e.g., different lineages of cells during mouse embryonic development, as we combine the early-, mid-, and late-blastocyst cells to gain a large enough sample size. While it is possible that this might lead to false positives in identifying coordinated bursting events, it will result in a decrease in power for the testing of differential bursting kinetics. Given the amount of stochasticity that is observed in the allele-specific expression data, how to define cell sub-types and how to quantify between-cell heterogeneity need further investigation.

## Conclusions

We have developed SCALE, a statistical framework for systematic characterization of ASE using data generated from scRNA-seq experiments. Our approach allows us to profile allele-specific bursting kinetics while accounting for technical variability and cell size difference. For genes that are classified as biallelic bursty through a Bayes categorization framework, we further examine whether transcription of the paternal and maternal alleles are independent, and whether there are any kinetic differences, as represented by bursty frequency and burst size, between the two alleles. Our results on the re-analysis of Deng et al. [2] and Borel et al. [17] provide insights into the extent of differences, coordination, and repulsion between alleles in transcriptional bursting.

## Methods

### Input for endogenous RNAs and exogenous spike-ins

For endogenous RNAs, SCALE takes as input the observed allele-specific read counts at heterozygous locus 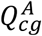 and 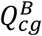, with adjustment by library size factor:

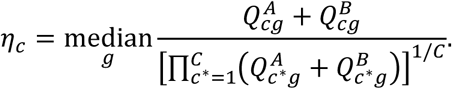

In addition, for spike-ins, SCALE takes as input the true concentrations of the spike-in molecules, the lengths of the molecules, as well as the depths of coverage for each spike-in sequence across all cells (Table S1A-S1C). The true concentration of each spike-in molecule is calculated according to the known concentration (denoted as *C* attomoles/uL) and the dilution factor (x40000):

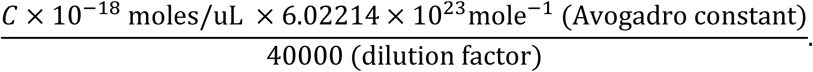

The observed number of reads for each spike-in is calculated by adjusting for the library size factor, the read length, and the length of the spike-in RNA. The bioinformatic pipeline to generate the input for SCALE is included in Supplementary Methods.

### Empirical Bayes method for gene categorization

We propose an empirical Bayes method that categorizes gene expressions across cells into silent, monoallelic, biallelic states based on their ASE data. Without loss of generality, we focus on one gene here with the goal of determining the most likely gene category based on its ASE pattern. Let 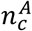 and 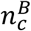 be the allele-specific read counts in cell *c* for allele A and B, respectively. For *each* cell, there are four different categories based on its ASE – {*∅, A, B, AB*} corresponding to scenarios where both alleles are off, only A allele is expressed, only B allele is expressed, and both alleles are expressed, respectively. Let *k* ∈ {1, 2, 3, 4} represent this cell-specific category. The log-likelihood for the gene across all cells can be written as: 

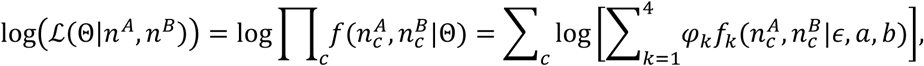

where the parameters are Θ = {*φ*_1_,…,*ϵ*_4_, *∈, a, b*} with 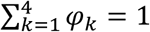 and each *f_k_* is a density function parameterized by *ϵ*, *a*, *b*. *ϵ* is the per-base sequencing error rate, and *a* and *b* are hyper-parameters for a Beta distribution, where *θ*_c_~Beta(*a, b*) corresponds to the relative expression of A allele when both alleles are expressed. It is easy to show that

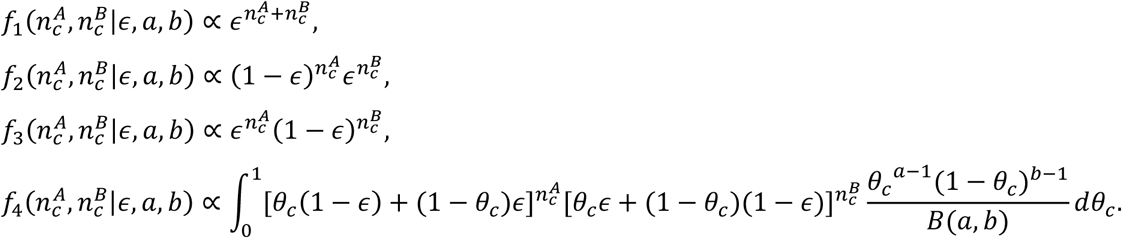

*ϵ* can be estimated using sex chromosome mismatching or be prefixed at the default value, 0.001. We require *a* = *b* ≥ 3 in the prior on θ_c_ so that the AB state is distinguishable from the A and B states. This is a reasonable assumption in that most genes have balanced ASE on average and the use of Beta distribution allows variability of allelic ratio across cells. We adopt an EM algorithm for estimation, with *Z* being the missing variables:

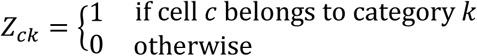

The complete-data log-likelihood is given as

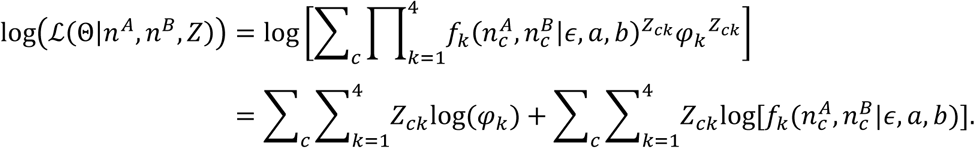

For each cell, we assign the state that has the maximum posterior probability and only keep a cell if its maximum posterior probability is greater than 0.8. Let *N*_∅_, *N_A_, N_B_* and *N_AB_* be the number of cells in state {∅}, {*A*}, {*B*}, and {*AB*}, respectively. We then assign a gene to be: (i) silent if *N_A_* = *N_B_* = *N_AB_* = 0 (ii) A-allele monoallelic if *N_A_* > 0, *N_B_* = *N_AB_* = 0; (iii) B-allele monoallelic if *N_B_* > 0, *N_A_* = *N_AB_* = 0 (iv) biallelic otherwise (biallelic bursty if 0.05 ≤ (*N_A_* + *N_AB_*)/(*N_ϕ_* + *N_A_* + *N_B_* + *N_AB_*) ≤ 0.95 and 0.05 ≤ (*N_B_* + *N_AB_*)/(*N_ϕ_* + *N_A_* + *N_B_* + *N_AB_*) ≤ 0.95).

### Parameter estimation for Poisson-Beta hierarchical model

Since exogenous spike-ins are added in a fixed amount and don’t undergo transcriptional bursting, they can be used to directly estimate the technical-variability-associated parameters {*α*,*β*, *κ*,*τ*} that are shared across all cells from the same sequencing batch. Specifically, we use non-zero read counts to estimate *α* and *β* through log-linear regression:

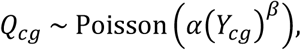

where *Q*_*cg*_ > 0, capture and sequencing efficiencies are confounded in *α* and amplification bias is modeled by *β* (Figure S5). We then use the Nelder-Mead simplex algorithm to jointly optimize *κ* and *τ*, which models the probability of non-dropout, using the likelihood function:

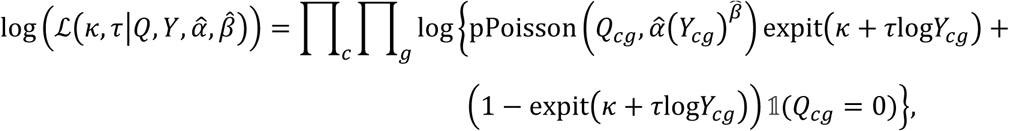

where pPoisson(*x, y*) specifies the Poisson likelihood of getting *x* from a Poisson distribution with mean *y*. This log-likelihood function together with the estimated parameters decomposes the zero read counts (*Q_cg_* = 0) into being from the dropout events or from being sampled as zero from the Poisson sampling during sequencing (Figure S5A).

The allele-specific kinetic parameters are estimated via the moment estimator methods, which is more computational efficient than the Gibbs sampler method adopted by Kim and Marioni [25]. For each gene, the distribution moments of the A allele given true expression levels 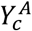 and 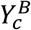 are:

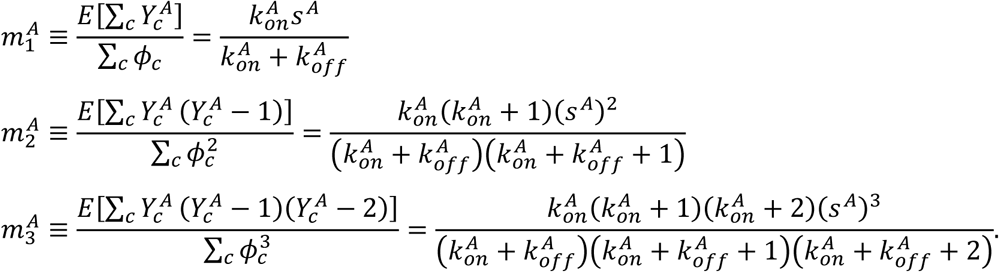

Solving this system of three equations, we have:

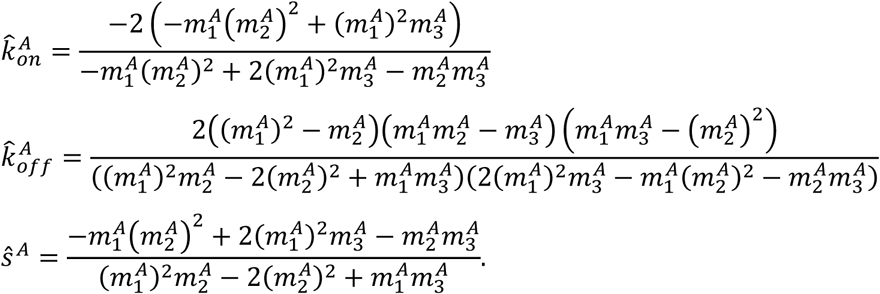

Substituting A with B we get the kinetic parameters for the B allele. To get the sample moments, we propose a novel histogram repiling method that gives the sample distribution and sample moment estimates of the true expression from the distribution of the observed expression (Figure S7). Specifically, for each gene we denote *c(Q)* as the number of cells with observed expression *Q* and *n(Y)* as the number of cells with the corresponding true expression *Y*. *c(Q)* follows a Binomial distribution indexed at *n(Y)* with probability of no dropout:

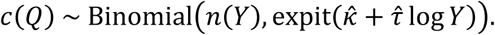

Then,

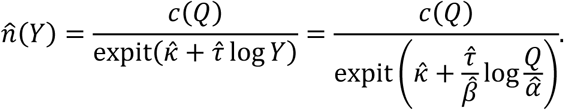

These moment estimates of the kinetic parameters are sometimes negative as is pointed out by Kim and Marioni [25]. By *in silico* simulation studies, we investigate the estimation accuracy and robustness under different settings.

### Hypothesis testing framework

We carry out a nonparametric bootstrap hypothesis testing procedure with the null hypothesis that the two alleles of a gene share the same kinetic parameters (Figure 4A-4B). The procedures are as follow.

i. For gene *g*, let 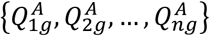 and 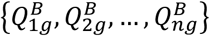 be the observed allele-specific read counts. Estimate allele-specific kinetic parameters with adjustment of technical variability:

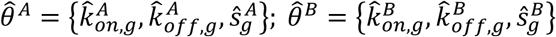
ii. Combine the 2*n* observed allelic measurements and draw samples of size 2*n* from the combined pool with replacement. Assign the first *n* with their corresponding cell sizes to allele A as 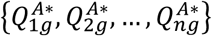 the next *n* to allele B 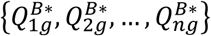 Estimate kinetic parameters with adjustment of technical variability from the bootstrap samples: 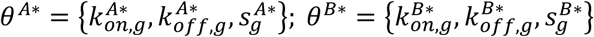 Iterate this times.
iii. Compute the *p*-values:

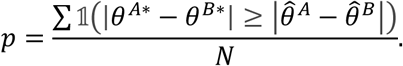

We adopt a Binomial test of allelic imbalance with the null hypothesis that the allelic ratio of the mean expression across all cells is 0.5. Chi-square test of independence is further performed to test whether the two alleles of a gene fire independently (Figure 4C). The observed number of cells is from the direct output of the Bayes gene categorization framework. For all hypothesis testing, we adopt FDR to adjust for multiple comparisons.

## Abbreviations

scRNA-seq: single-cell RNA sequencing
ASE: allele-specific expression
SNP: single-nucleotide polymorphism
RNA-seq: RNA sequencing
ME: monoallelic expression
RME: random monoallelic expression
FISH: fluorescence *in situ* hybridization
EM: expectation-maximization
FDR: false discovery rate
RPKM: reads per kilo base per million reads
PCA: principal component analysis
QC: quality control
QTL: quantitative trait loci

## Declarations

We thank Dr. Daniel Ramsköld for providing dataset of the mouse preimplantation embryos, Dr. Christelle Borel for providing dataset of the human fibroblast, and Cheng Jia, Dr. Arjun Raj, and Dr. Uschi Symmons for helpful comments and suggestions. This work was supported by National Institutes of Health (NIH) grant R01HG006137 to NRZ, and R01GM108600 and R01HL113147 to ML. The authors declare that they have no competing interests.

